# A modular neural network model of grasp movement generation

**DOI:** 10.1101/742189

**Authors:** Jonathan A. Michaels, Stefan Schaffelhofer, Andres Agudelo-Toro, Hansjörg Scherberger

## Abstract

One of the primary ways we interact with the world is using our hands. In macaques, the circuit spanning the anterior intraparietal area, the hand area of the ventral premotor cortex, and the primary motor cortex is necessary for transforming visual information into grasping movements. We hypothesized that a recurrent neural network mimicking the multi-area structure of the anatomical circuit and using visual features to generate the required muscle dynamics to grasp objects would explain the neural and computational basis of the grasping circuit. Modular networks with object feature input and sparse inter-module connectivity outperformed other models at explaining neural data and the inter-area relationships present in the biological circuit, despite the absence of neural data during network training. Network dynamics were governed by simple rules, and targeted lesioning of modules produced deficits similar to those observed in lesion studies, providing a potential explanation for how grasping movements are generated.

## Introduction

Interacting with objects is an essential part of daily life for primates. Grasping is one of our most complex behaviors, requiring the determination of object features and identity, followed by the execution of the correct temporal sequence of precise muscle patterns in the arm and hand necessary to reach and grasp the object. In macaque monkeys, the circuit formed by the anterior intraparietal area (AIP), the hand area (F5) of the ventral premotor cortex, and the hand area of the primary motor cortex (M1) is essential for grasping. These areas share extensive anatomical connections (Luppino et al., 1999), forming a long-range circuit (Figure 1a) where AIP receives the largest amount of visual information, and M1 has the largest output to the brainstem and spinal cord. All three areas have been shown to contain grasp-relevant information well before movement (Baumann et al., 2009; Carpaneto et al., 2011; Fluet et al., 2010; Murata et al., 1997, 2000; Schaffelhofer et al., 2015a).

**Fig. 1.**
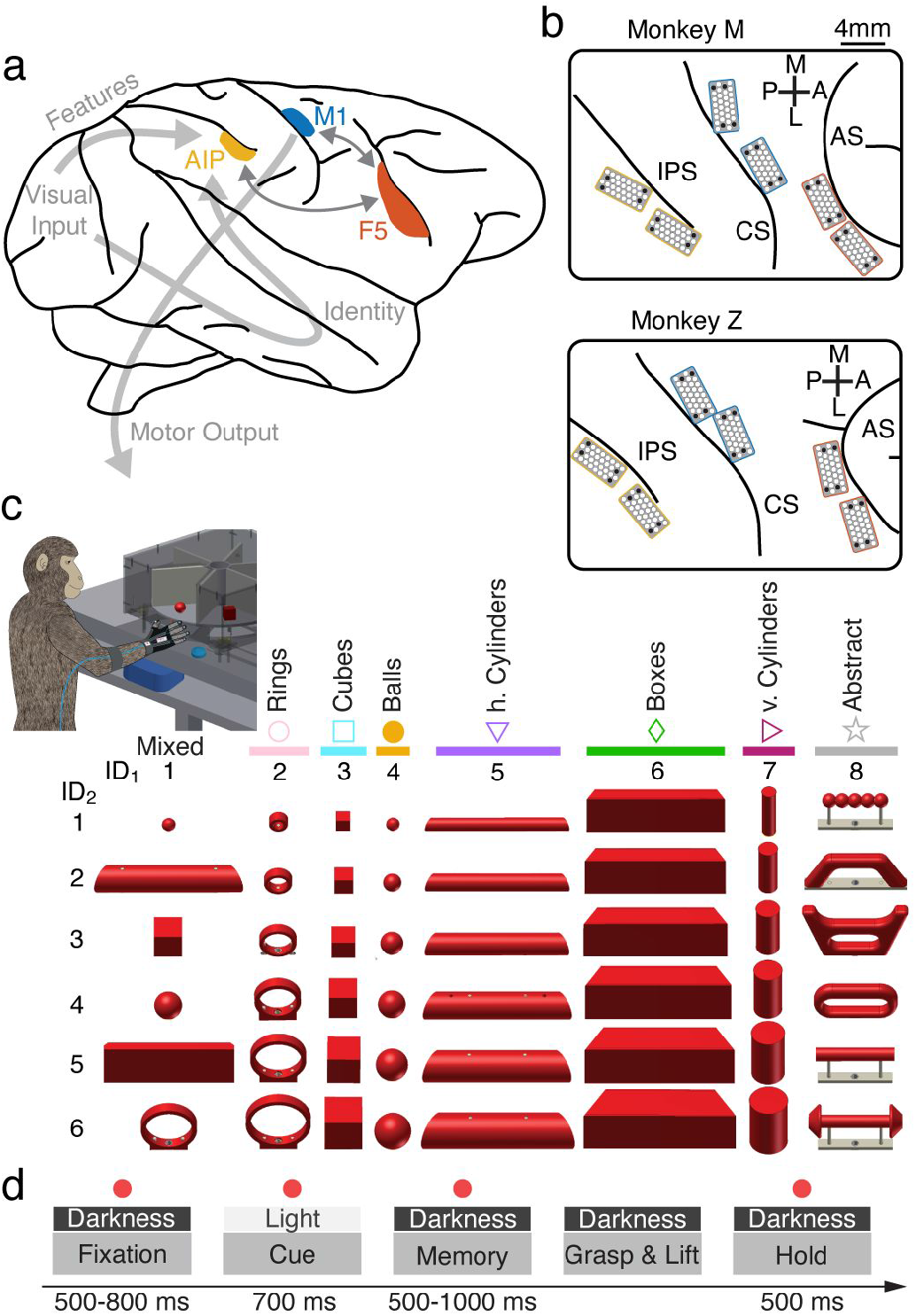
Fronto-parietal grasping circuit and experimental design. (a) Simplified brain schematic of the fronto-parietal grasping circuit. Visual information is processed in two parallel streams carrying primarily object features or identity information, both converging on the anterior intraparietal sulcus (AIP). AIP has strong reciprocal connections with the hand area (F5) of the ventral premotor cortex, which has strong reciprocal connections to the hand area of the primary motor cortex (M1). M1 has the majority of subcortical and spinal cord output projections. (b) Location of implanted floating micro-electrode arrays, covering the three desired regions. Arcuate sulcus (AS), central sulcus (CS), intraparietal sulcus (IPS), anterior (A), posterior (P), medial (M), lateral (L). Black dots represent ground and reference electrodes. (c) Monkeys sat in front of a motorized turntable that presented one of six objects to be grasped on any given trial (reproduced from (Schaffelhofer and Scherberger, 2016)). Multiple turntables presented in random order across sessions allowed for a total of 42 objects. Gloves with magnetic sensors allowed full tracking of arm and hand kinematics on single trials. (d) Trials began with visual fixation of a red dot for a variable period. Objects were illuminated temporarily, and monkeys were required to withhold movement until a go cue (blinking of fixation dot) instructed them to grasp and lift the object in darkness. Eye fixation was enforced throughout each trial.

Reversible inactivation of AIP (Gallese et al., 1994; Tunik et al., 2005) or F5 (Fogassi et al., 2001) results in a selective deficit in appropriately pre-shaping the hand during grasping, while M1 lesions lead to profound hand movement deficits (Hoffman and Strick, 1995; Murata et al., 2008; Passingham et al., 1983), providing evidence that these areas are required for successful grasping. Additionally, M1 has the largest density of projections directly onto motor neurons for control of the fingers, and precise finger control does not normally recover after lesion (Murata et al., 2008; Passingham et al., 1983). So far, models of the grasping system have relied on manually tuning the properties of individual neurons to match the assumed role of a given region (Fagg and Arbib, 1998). No comprehensive model exists of the entire transformation between vision and action, limiting our ability to understand the flexibility of the grasping system.

Goal-driven modeling has emerged as a powerful tool for generating potential neural mechanisms explaining various behaviors (Yamins and DiCarlo, 2016). The creation of vast datasets of labeled images (Imagenet, Deng et al., 2009) opened the door to studying the computational principles underlying object identification using convolutional neural networks (CNNs), such as Alexnet (Krizhevsky et al., 2012). Feedforward modeling of the ventral stream using CNNs has led to powerful insights into hierarchy of brain networks (Cadieu et al., 2014; Yamins et al., 2014), revealing that subsequent layers of CNNs for object identification align well with brain regions along the ventral stream. Similar approaches have been used in retinal modeling (Maheswaranathan et al., 2018), and recent studies incorporating recurrence into CNNs (Kar et al., 2019; Kietzmann et al., 2019; Nayebi et al., 2018). In parallel, advances have been made in understanding motor cortex by modeling it as a dynamical system (Churchland et al., 2012; Shenoy et al., 2013) implemented as a recurrent neural network (RNN) (Hennequin et al., 2014; Michaels et al., 2016; Stroud et al., 2018; Sussillo et al., 2015). In these models, and likely in motor cortex (Churchland et al., 2012), preparatory activity sets initial conditions that unfold predictably to control muscles during reaching.

In the current work, we bridge the gap between previous work in visual processing and motor control by modeling the entire processing pipeline from visual input to muscle control of the arm and hand. Firstly, we recorded neural activity from AIP, F5, and M1 of two macaque monkeys while they grasped a diverse set of 42 objects. Activity in AIP was best explained by visual features extracted from penultimate layers of VGG (Simonyan and Zisserman, 2014), a CNN trained to identify objects. M1 activity was best explained by muscle kinematics (i.e. muscle fiber velocity), and F5 was intermediate between the two. Based on these results, we devised a number of neural network architectures to model the function of this circuit. Primarily, we trained an mRNN with sparsely connected modules mimicking cortical areas to use visual features from VGG to produce the muscle kinematics required for grasping. Additionally, we trained networks with varying degrees of sparsity between modules, only feedforward connections between modules, homogeneous or sparse connectivity lacking modules, and others.

While all architectures were able to produce the required muscle kinematics, they differed in their ability to explain the neural population dynamics observed in the brain. Specifically, modular networks with relatively sparse inter-module connectivity (10%) in both the feedforward and feedback direction that received visual features extracted from later layers of the CNN matched neural data better than all other models, suggesting that these may be organizational principles of the cortical circuit. Fixed-point analysis revealed that networks used a simple strategy to complete the task, reorganizing activity during the cue and memory period that was released during movement to generate the appropriate muscle kinematics, suggesting a flexible strategy that might generalize well to new input. Interestingly, targeted lesioning of modules specifically in mRNN models that penalized high firing rates produced behavioral deficits that varied by module and paralleled deficits observed in previous lesion studies of these cortical areas, providing a potential explanation of how these cortical regions may complete this task in tandem.

## Results

### Kinematic and neural activity recorded during a many-object grasping task

To investigate the computational basis of grasp control, we recorded neural activity in the inter-connected anterior intraparietal area (AIP), the hand area (F5) of ventral premotor cortex, and the hand area of motor cortex (M1) using floating micro-electrode arrays (Fig. 1a,b) while two rhesus macaques (monkeys M and Z) performed a delayed grasping task. We presented monkeys with 42 objects composed of shapes of various sizes and orientations on a series of rotating turntables (Fig. 1c). Turntables were presented in random order each session. Experimental and behavioral findings have been presented in previous works (Schaffelhofer and Scherberger, 2016; Schaffelhofer et al., 2015a). Monkeys wore a glove that allowed full joint tracking (Schaffelhofer and Scherberger, 2012) of the arm, hand, and fingers on single trials, and this data was further transformed into muscle space using a previously described musculoskeletal model (Delp et al., 2007). On individual trials monkeys had to fixate a red point just under each object, after which the object was illuminated temporarily. Monkeys then waited for a go cue in darkness, after which they reached to, grasped, and lifted the object (Fig. 1d).

We analyzed the data from 10 recording sessions per monkey (labeled M1-M10 and Z1-Z10). On average, each recording session of monkey M consisted of 549±35 (Mean±S.D.) trials, and 153±8, 179±7, and 215±14 single- and multi-units were recorded from AIP, F5, and M1, respectively. On average, each recording session of monkey Z consisted of 490±25 (Mean±S.D.) trials, and 122±10, 137±6, and 126±9 single- and multi-units were recorded from AIP, F5, and M1, respectively.

Previous work using this dataset (Schaffelhofer and Scherberger, 2016; Schaffelhofer et al., 2015a) has shown that this circuit is heavily modulated by the type of object being grasped, containing rich information about objects, both during the presentation and the intervening memory period, and representing temporal information about the kinematic signals required for grasping during movement. Next, we wanted to determine how visual information about grasp targets is used and transformed into the information necessary to execute grasping.

### Graded shift from visual to kinematic features in the grasping circuit

The first step in designing a model of the grasping circuit was determining what information may be available during the planning of grasping movements. Based on the established role of AIP in grasping and its connectivity to areas containing information about the size, shape, orientation, and identity of objects, we hypothesized that later layers of existing convolutional neural network (CNN) models of the ventral stream may match activity in AIP. We constructed simulated images of the task from the monkey’s perspective (Fig. S1) and fed them into VGG (Simonyan and Zisserman, 2014) (Fig. 2a), a feedforward CNN that was pre-trained to identify objects in ImageNet (Deng et al., 2009), which contains millions of images. We read out the hidden activity from each of the network layers and compared their ability to explain neural activity in each brain area during the early cue period, when the object was visible, using a single-trial, cross-validated regression method very similar to an established metric in the visual system (Methods, “BrainScore” Schrimpf et al., 2018).

**Fig. 2.**
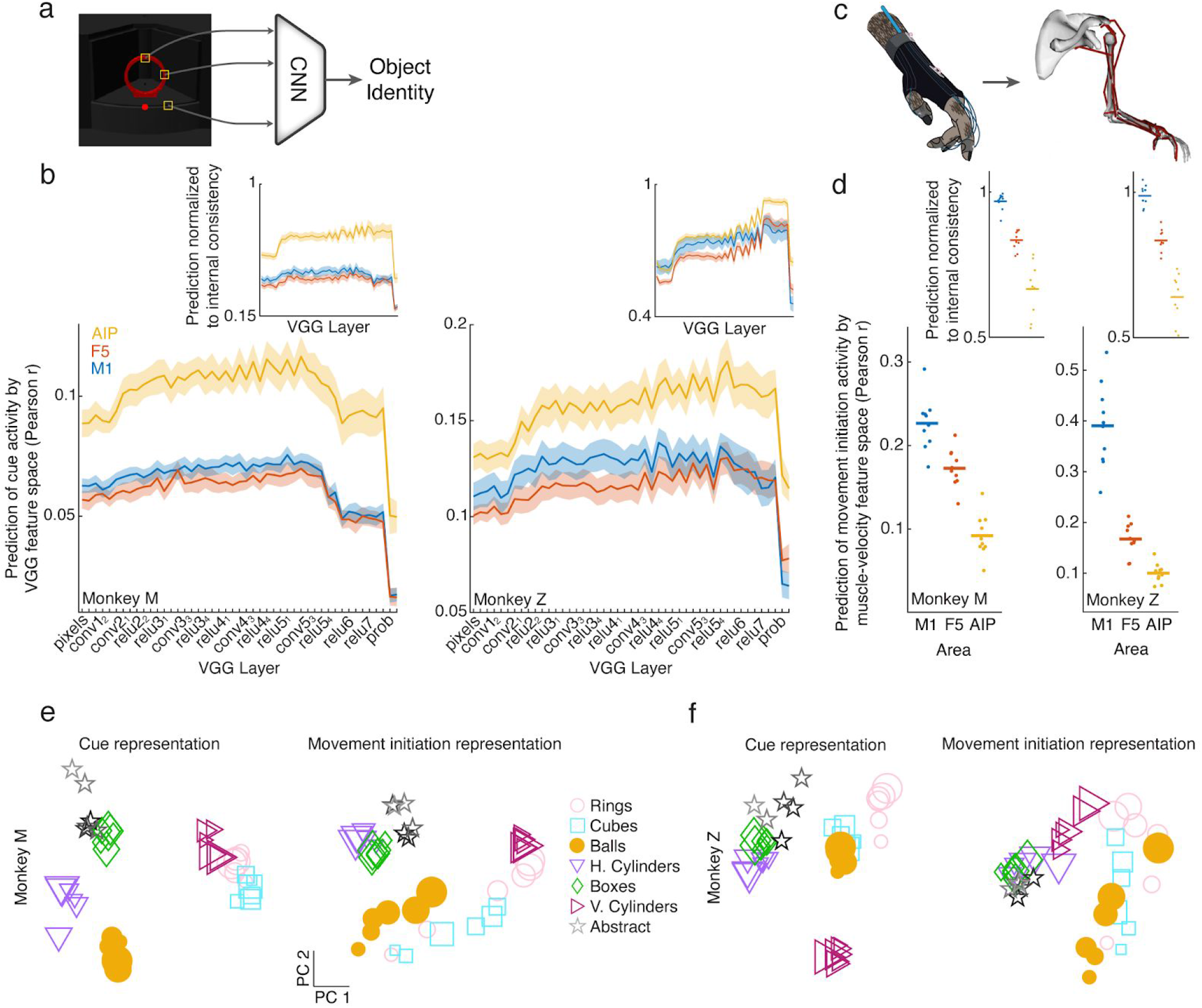
Graded shift from visual to kinematic features in the fronto-parietal grasping circuit. (a) Simulated images of all objects were fed through a convolutional neural network (CNN) pre-trained to extract object identity (VGG). (b) The representation of all objects in each layer of the CNN (first 20 principal components) was regressed against the single-trial neural activity of each unit during the early portion of the cue period, and the median fit was taken over all units within one recording session. Solid line and error surfaces represent the mean and s.e.m. over all recording sessions of each monkey. (b - inset) To ensure that results were not due to varying signal quality or firing rate between areas, insets show regression results normalized to internal consistency of each area (i.e. half of trials correlated with the other half condition-wise; Methods). (c) Joint angles (27 DOF) were transformed into muscle length space (50 DOF) using a musculoskeletal model. For visualization purposes, not all muscles are shown. (d) The mean muscle velocity of all grasping conditions during movement initiation (200 ms before to 200 ms after movement onset) was regressed against the single-trial neural activity of each unit during the same time period. Each point represents one recording session of each monkey (monkey M - left, monkey Z - right). (d - inset) Same normalization procedure as in (b - inset). (e) Example neural representation (first two principal components) of each object across all three areas during the cue period and during movement initiation (session M7). The size of each marker indicates the relative size of each grasping object. (f) Same as (e), but for session Z9.

As predicted, VGG features better explained single-trial activity in AIP than in F5 or M1 (Fig. 2b, monkey M - F5 and M1, paired t-test p = 2e-200, 6e-143; monkey Z - F5 and M1, paired t-test p = 6e-99, 4e-61). Additionally, the later layers of VGG (e.g. relu5_4 layer) predicted activity in AIP better than the early layers (pixel layer, monkey M - paired t-test p = 0.01, monkey Z - paired t-test = 0.0001), suggesting that VGG produced features that were more predictive of neural activity in AIP than pure pixel information.

To control for differences in firing rate and recording quality between areas, we calculated the internal consistency of each area (i.e. how well single-trial responses correlate across repetitions) and normalized to that value (Methods). A normalized value around 1 would indicate that a set of predictors captures the condition-dependent neural features as well as can be expected given the reliability of the recorded data. In both monkeys, the above results remained unchanged (Fig. 2b inset, monkey M - F5 and M1, paired t-test p = 3e-228, 7e-95; monkey Z - F5 and M1, paired t-test p = 7e-95, 7e-21; relu5_4 vs. pixel layer, monkey M - paired t-test p = 4e-5; monkey Z - paired t-test p = 6e-10). Furthermore, the advanced layers of VGG achieved a normalized fit between 0.7-0.9 in AIP, suggesting that those visual features well explain neural activity in AIP during the cue period. Very similar results were obtained by feeding our images through both Alexnet (Krizhevsky et al., 2012) and Resnet (He et al., 2015), two widely used CNN architectures. Similar results are also obtained using the later portion of the cue period (400-700ms after onset), although these results were more homogenous between areas, suggesting a strong influence of recurrence between regions.

Having established that later layers of a CNN trained to identify objects provide natural inputs to AIP, the next step was to determine reasonable outputs of the grasping circuit. As mentioned previously, monkeys wore a tracking glove (Schaffelhofer and Scherberger, 2012) that allowed the extraction of 27 degrees of freedom of movement information, almost completely capturing reach to grasp movement trajectories. The joint angle signal was further transformed into a 50-dimensional muscle space using a musculoskeletal model of the primate arm and hand (Schaffelhofer et al., 2015b) (Fig. 2c), allowing detailed access to muscle kinematics in the hand that would be very difficult to obtain using single muscle recording techniques. While this model does not give us direct access to muscle force or activity, it provides a kinematic signal that bears many similarities to muscle activity, especially during the early portion of the movement before the hand makes contact with the object. We opted to analyze the muscle velocity signal, since it is invariant to starting hand posture. Similar to the analysis of visual features, we used a 50-dimensional muscle velocity signal to predict the activity of individual units around movement onset (200 ms before to 200 ms after movement onset). Muscle features better predicted activity in M1 than F5 or AIP (Fig. 2d, monkey M - F5 and AIP, paired t-test p = 0.008, 2e-6; monkey Z - F5 and AIP, paired t-test p = 4e-5, 7e-7), showing the opposite correspondance with cortical regions as the visual feature analysis. Furthermore, the results did not change when controlling for internal consistency (Fig. 2d inset, monkey M - F5 and AIP, paired t-test p = 2e-5, 3e-6; monkey Z - F5 and AIP, paired t-test p = 5e-7, 2e-7), with normalized fits around 1 for both monkeys, suggesting that M1 data is predicted as well as possible by the muscle velocity signal.

Together, these results strongly suggest a visuomotor gradient from AIP to F5 to M1 that transforms visual features of objects into muscle kinematic signals. However, these analyses only provide snapshots in time and cannot explain the temporal evolution of neural population activity nor the computational mechanisms required to complete the task. In Figure 2e-f, we visualize the relationship between different objects and sizes of objects in the two neural dimensions of largest variability, showing how the relationship between objects is reorganized between the cue and movement periods (see Fig. S2 for equivalent visualization of all CNN layers). For example, while size does not seem to play a large role in the cue period, some objects are organized by size during movement initiation in both monkeys (balls, cubes, rings). One of the goals of the current work is to provide potential explanations of *how* such reorganization may take place, and *why* this representation is useful for movement generation.

### A modular recurrent neural network model of vision to hand action

To build a comprehensive model of the grasping circuit incorporating temporal dynamics, we devised a modular recurrent neural network (mRNN) inspired by the above results and the known anatomical connectivity of the grasping circuit (Methods). The model consisted of three interconnected stages designed to reproduce the muscle dynamics necessary to grasp objects (Fig. 3a). The visual input was a 20-dimensional visual feature signal consisting of the first 20 principal components of the features in one of the layers of VGG (relu5_4) that was a good match to AIP activity while viewing the simulated images. This visual signal entered the input module, a fully-connected RNN (all modules used a saturating nonlinearity, the rectified hyperbolic tangent, ReTanh), that relayed information to the intermediate module through sparse connectivity (10%). Similarly, the intermediate module projected to the output module sparsely, and equally sparse feedback connections existed for each of the feedforward connections. In order to match kinematic timing, all three modules received a hold signal that cued movements 200 ms before desired movement onset, which was approximately when the monkey’s hand lifted off of a handrest button. The output module was most directly responsible for generating the 50-dimensional muscle velocity signal required to grasp each object up to 400 ms into movement and to suppress movement earlier in the trial. Figure 3b shows inputs for an example trial, including the visual cue signal and the hold signal. During the fixation, memory, and movement periods only the fixation point was presented, while during the go cue the fixation point disappeared for 100 ms.

**Fig. 3.**
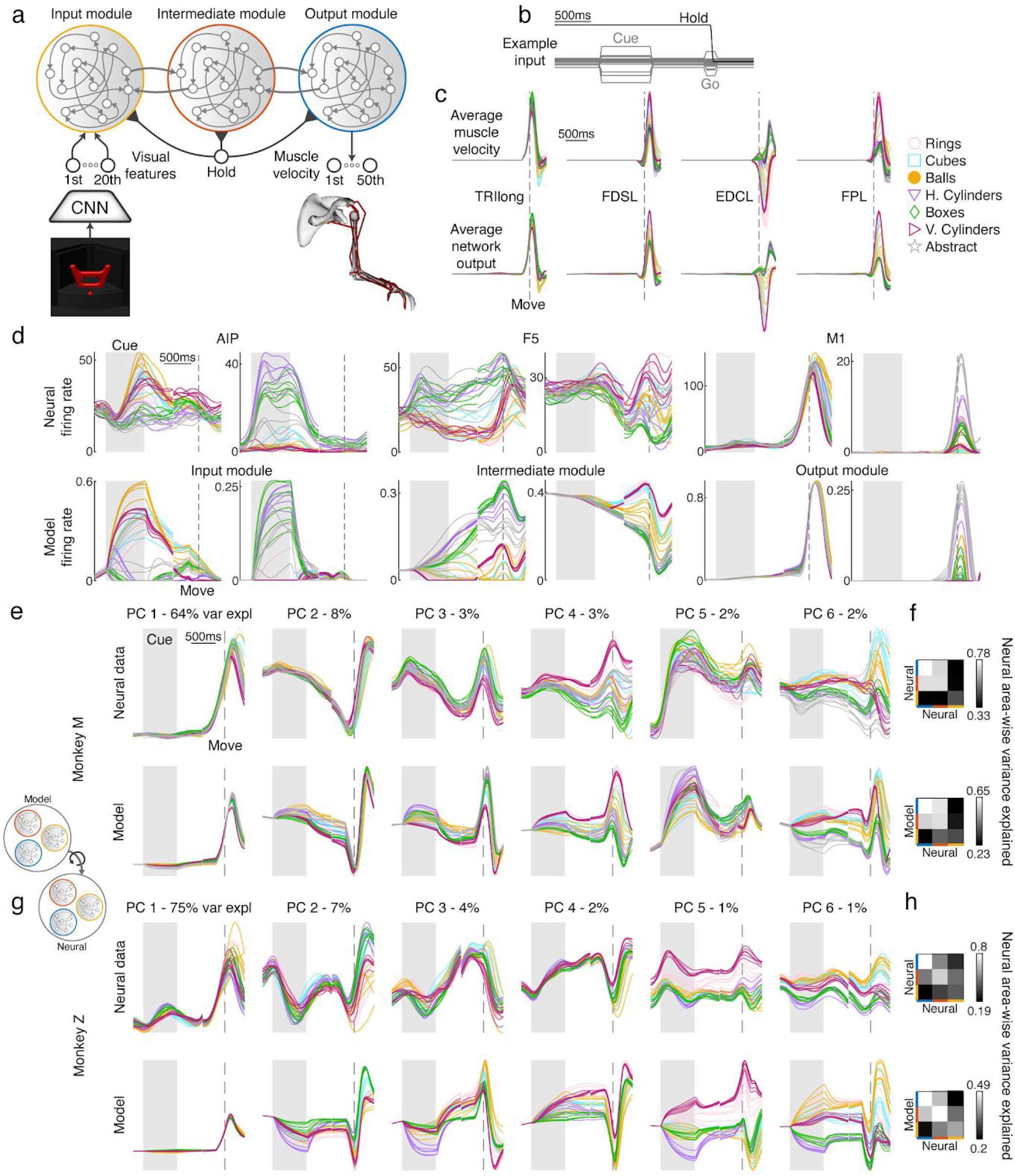
Modular recurrent neural network model of the fronto-parietal grasping circuit. (a) Schematic of neural network model. Visual features of each object (first 20 principal components of relu5_4 layer) are fed into an input module, which is sparsely reciprocally connected to an intermediate module, which is similarly connected to an output module. The output module must recapitulate the muscle velocity for every object grasped by the monkey. Every module received a hold signal that is released 200 ms prior to movement onset. (b) Example input for an exemplary trial. (c) Average muscle velocity for four example muscles (TRIlong - triceps long head, FDSL - flexor digitorum superficialis digit 5, EDCL - extensor digitorum communis digit 5, FPL - flexor pollicis longus) showing exemplar recorded kinematics and network output (session M2). (d) Two example units from each pair of modules and brain regions showing similar properties and highlighting common features of each area. Traces were aligned to two events, cue onset and movement onset, and concatenated together. The shaded gray area represents the cue period, while the dashed line represents movement onset. (e) Procrustes analysis comparing the dynamics of an exemplar mRNN model (ReTanh activation function, rate regularization - 1e-1, weight regularization - 1e-5, inter-module sparsity - 0.1) with neural data recorded from AIP, F5, and M1 (session M2). Fits of the simulated to neural data were projected onto the first 6 PCs defined on the neural data for visualization purposes only, and percentages show variance explained in biological data per PC. (f) Area-wise procrustes was performed between each brain region and itself (top, Methods), or between each module and brain region (bottom). (g,h) same as e,f, but for session Z4. For c,d,e,g, the multiple traces for each type of object represent the different sizes within a turntable.

We used an optimization procedure (Hessian-free optimization, Methods) to train a series of networks to recapitulate the average movement behavior of each monkey, while also varying many aspects of the network architecture or regularization (total of 1760 trained networks, architectures detailed in later sections). Each network was trained to reproduce the average muscle velocity of each condition using a random set of two trials (42 conditions × 2 examples) where the timing parameters of each example was drawn randomly from the set of timing parameters observed during the monkey experiments (Figure 1d). It is crucial to emphasize that no neural data was used in any training procedure, allowing us to compare the neural dynamics of the recorded data to the internal dynamics of our model. Trained networks were very successful in reproducing the desired muscle kinematics (Fig. 3c), achieving low levels of normalized error (average of 6% for unregularized networks). In addition to successful recapitulation of muscle kinematics, networks were also able to suppress output before the movement period and maintain an internal representation of the task conditions in the absence of a visual cue.

In addition to the task goal we tested the effect of common constraints during training via two regularizations (Methods): 1) a cost on the firing rate of all neurons (L2 rate), and 2) a cost in the input and output weights of the network (L2 weight). Hyper-parameters for each regularization were tested systematically in a later analysis, and an exemplar network with an L2 rate regularization of 1e-1 and an L2 weight regularization of 1e-5 is analyzed in the following section.

### mRNN model with visual feature input reproduces single unit, population level, and area-wise neural dynamics

To gain an initial intuition of how the hidden state of the regularized mRNN compared to neural data, we plotted the average firing rates of 6 example units that showcase the similarities between the modules and the brain regions of interest (Fig. 3d). Units in AIP and the input module were often characterized by large responses to the visual cue that were either partially maintained through the memory period into movement, or decayed rapidly after the disappearance of the stimulus. Units in F5 and the intermediate module often showed sustained responses throughout the trial that were sensitive to time within the trial. M1 and output module units showed the largest response during movement itself, but often had stable or ramping activity earlier in the trial.

While these example units are useful insights into both the simulation and the neural data, a proper characterization requires a full analysis of the neural population dynamics. To capture similarities between the population dynamics of neural and simulated data, we devised a number of metrics based on procrustes analysis (Schönemann, 1966). Imagine we have two shapes in front of us, for example a square and a triangle, consisting of the set of two dimensional points that make up each shape. If we would like to see if it’s possible to overlap the triangle on the square *without distorting the overall shapes*, procrustes analysis provides a method for finding the optimal rotation that aligns them in arbitrary dimensionality (Methods). This method is ideal for comparing simulated and biological data, where the square represents the neural activity of simulated data and the triangle biological data. Procrustes does not distort the variance structure of either set of data, and similar methods based on dot product similarity have been shown to be ideal for comparing artificial neural networks (Gretton et al., 2005; Kornblith et al., 2019). Other commonly used methods, such as canonical correlation analysis (CCA), distort the amount of variance explained by individual units, although this can be alleviated somewhat by first performing principal component analysis (PCA) to constrain the analysis to dimensions of highest variance (Maheswaranathan et al., 2019; Raghu et al., 2017; Sussillo et al., 2015).

The first of these analyses is shown in Figure 3e,g, comparing the dynamics of an exemplar mRNN to neural data across all brain regions. After rotating simulated data onto neural data, results were projected onto the first 6 PCs defined by the covariance of the neural data for visualization purposes. For both monkeys there was a striking similarity between the simulations and the empirical data, both in terms of temporal properties throughout the trial and how the different grasp conditions were organized. Temporal features were the most dominant, similar to previous work (Kaufman et al., 2016), while the more condition dependant signals were captured by dimensions of relatively small variance. Across all neurons the exemplar mRNN network was able to explain 65% of the variance in neural data averaged across recording sessions.

In addition to the overall ability of the model to explain the neural data across brain regions, we were interested in two additional metrics: 1) how well each module explains the brain region it was expected to mimic, and 2) how well the pair-wise fit between each module and each brain region matches the expected pairwise relationship in the biological circuit. For the first additional metric, we simply averaged the fit of each of the three modules to each corresponding brain region, explaining 54% of the variance in the exemplar mRNN (see Fig. S3 for example visualization). For the second additional metric, we computed the pair-wise fit of each module to each brain region using procrustes, and correlated those values with an estimate of the expected fit between brain regions using a resampling procedure (Methods), yielding a high correlation of 0.91 for the exemplar mRNN. Importantly, visualizing the pair-wise fits (Fig. 3f,h) shows that the modules within the model best explained the brain regions they were predicted to mimic, even though no neural data was used during training, suggesting that the task combined with the architecture of the model was able to reproduce inter-area differences observed in the grasping circuit.

We found that a modular network with a saturating nonlinearity (ReTanh), rate and weight regularization, and 10% connectivity between modules explained the largest amount of variance in the neural circuit. To explore how our choice of network parameters in the exemplar network affected these results, we systematically tested (Fig. 4a) the effect of activation function, rate regularization, weight regularization, and inter-module sparsity on the three metrics presented above (Fig. 4b-d), additionally generating and training 5 networks for each choice of hyper-parameters and averaging across the five repetitions. The results of these analyses showed small effects of activation function (ReTanh best, in contrast to (Yang et al., 2019)), rate regularization (1e-2 best), and weight regularization (no regularization best) across metrics. However, the sparsity between modules had a larger effect on the correlation in inter-area structure (Fig 4e), showing a poor fit when the modules were fully connected, and the best fit when modules were 10% connected, suggesting that an intermediate level of sparsity was required to properly model the inter-area differences, similar to what is observed in the biological circuit (Markov et al., 2014). In many cases, the highest level of weight regularization increased error significantly on the task (Fig. S4), so these networks were excluded from further analysis. Two example networks illustrate how the fits to neural data were not as good for ReLu networks with no sparisty between modules (Fig 4h,i) or ReTanh networks with 10% sparsity, but no regularization (Fig. 4j,k) as compared to our exemplar mRNN network (Fig. 3). Overall, this analysis suggests that sparsity between modules had the largest effect on the ability of various mRNN models to fit neural data. However, later analysis will show that predictions of regularized networks differ significantly from unregularized networks.

**Fig. 4.**
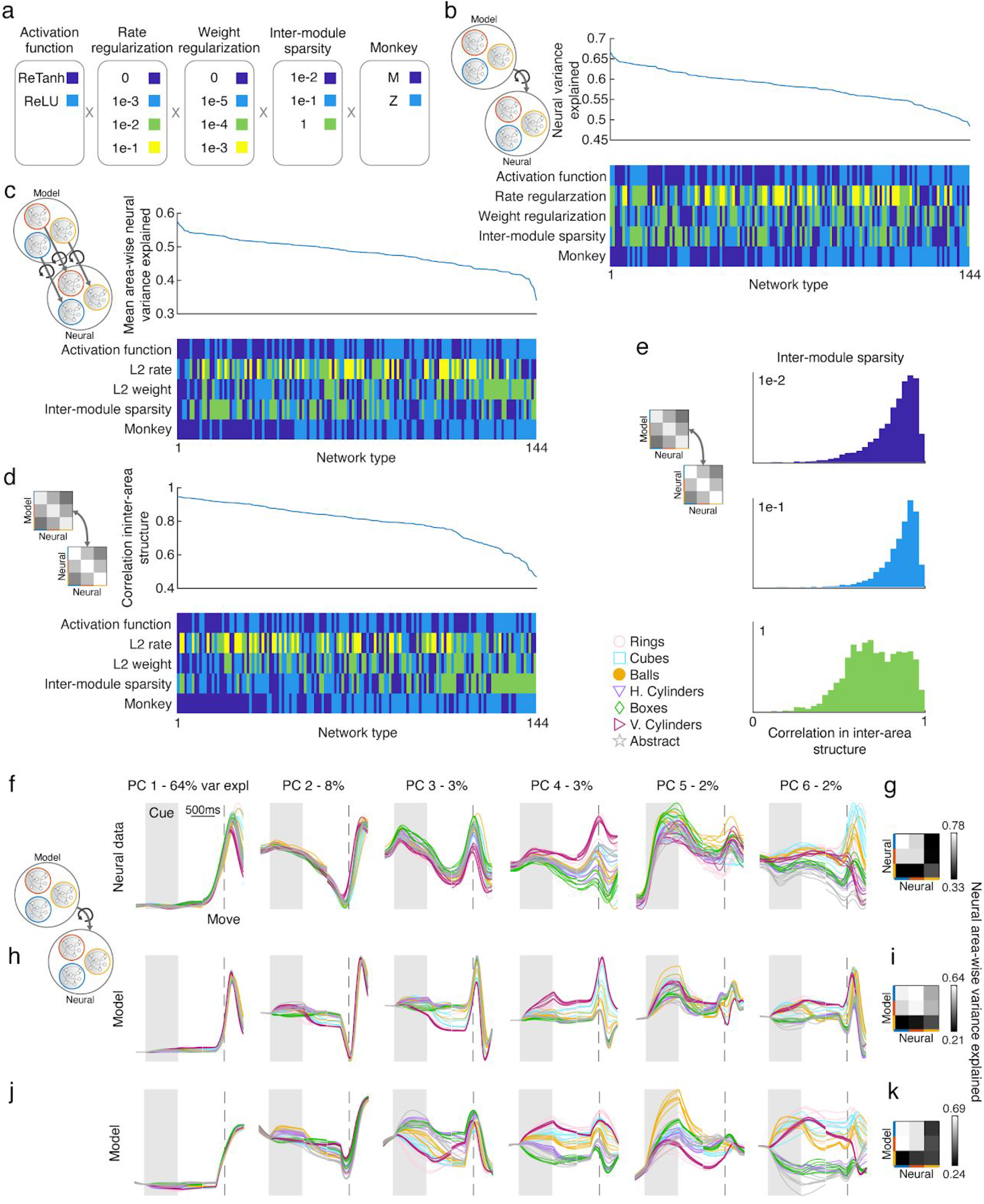
Effect of network parameters on the ability for the mRNN model to explain the biological circuit. (a) 720 networks were trained (5 repetitions of each network type), varying the activation function, rate and weight regularizations, inter-module sparsity, and the monkey being modeled. Note that the highest level of weight regularization was excluded, since task performance was severely affected. (b-d) Average performance across recording sessions for each of the three proposed metrics sorted from best to worst across all networks. (e) The effect of inter-module sparsity on the correlation of inter-area structure between the model and the predicted inter-area structure in the neural data. (f) Procrustes analysis comparing the dynamics of two exemplar mRNN models to neural data across all brain regions (session M2). Fits of the simulated to neural data were projected onto the first 6 PCs defined on the neural data for visualization purposes only, and percentages show variance explained in biological data per PC. (g) Area-wise procrustes was performed between each brain region and itself, or between each module and brain region (i,k). (h) Exemplar model with the parameters (ReLu activation function, L2 rate regularization - 1e-1, L2 weight regularization - 1e-5, inter-module sparsity - 1). (j) Exemplar model with the parameters (ReTanh activation function, rate regularization - 0, weight regularization - 0, inter-module sparsity - 0.1). For f,h,j, the multiple traces for each type of object represent the different sizes within a turntable.

### mRNN model with visual feature input outperforms alternative models

In the previous sections, we have shown that mRNN models with sparse inter-module connectivity, visual feature input from later layers of an object identification CNN, and trained to produce the muscle dynamics necessary for grasping are able to explain neural dynamics and inter-area differences across the AIP, F5, M1 grasping circuit. However, it was essential to test some alternative models to determine which of these design choices were most essential in producing this result. We tested 5 alternative models in addition to the Full mRNN model: 2) an mRNN model with only feedforward connections between modules (Feedforward) to test the necessity of top-down feedback; 3) an mRNN model receiving a labeled-line input, where each condition is represented by a separate input dimension (Labeled-Line) to see if equivalent visual inputs could develop by training on motor output alone; 4) an mRNN model with output conditions reassigned between grasping objects (Condition-Shuffled Output) to test if the precise matching between kinematics and neural data was necessary; 5) a fully-connected module (Homogeneous) to test if modular processing is necessary; or 6) a sparsely-connected module with input, hidden, and output sparsity matching the Full model (Sparse).

All of these alternative models were able to achieve task error similar to the Full model (average 5%) and were trained for the same set of rate and weight regularizations as previous models (Fig. 4). The results of the best set of regularizations across metrics (chosen separately for each architecture) are shown for each of the three previously introduced metrics in Figure 5a-c. Similar results are obtained if instead results are averaged across all sets of regularization parameters. Procrustes analysis across all brain regions revealed that all models performed more poorly than the Full model (Fig. 5a, paired t-test, p < 0.01), a result that was replicated when considering average area-wise fit (Fig. 5b). Comparing the correlation in inter-area structure between models showed that in general the same result held (Fig. 5c), although not significantly for the Feedforward and Labeled-Line networks in monkey Z. It’s important to note that we are not suggesting that a strictly 3-module network is necessary for explaining neural data in the grasping circuit, but rather that multi-module networks with distinct modules can explain neural data better than alternative models. Three was a natural choice based on the anatomical connectivity AIP, F5, and M1, as well as our access to recordings from all three regions. Two example networks illustrate how the fits to neural data were not as good for Homogeneous models (Fig 5f,g) or Condition-Shuffled Output models (Fig. 5h,i) as compared to our exemplar mRNN network (Fig. 3). Overall, this additional analysis reinforces that modular networks that receive visual feature input, have feedforward and feedback connections between modules, and produce the muscle kinematics of grasping provide the best explanation of neural data from the grasping circuit.

**Fig. 5.**
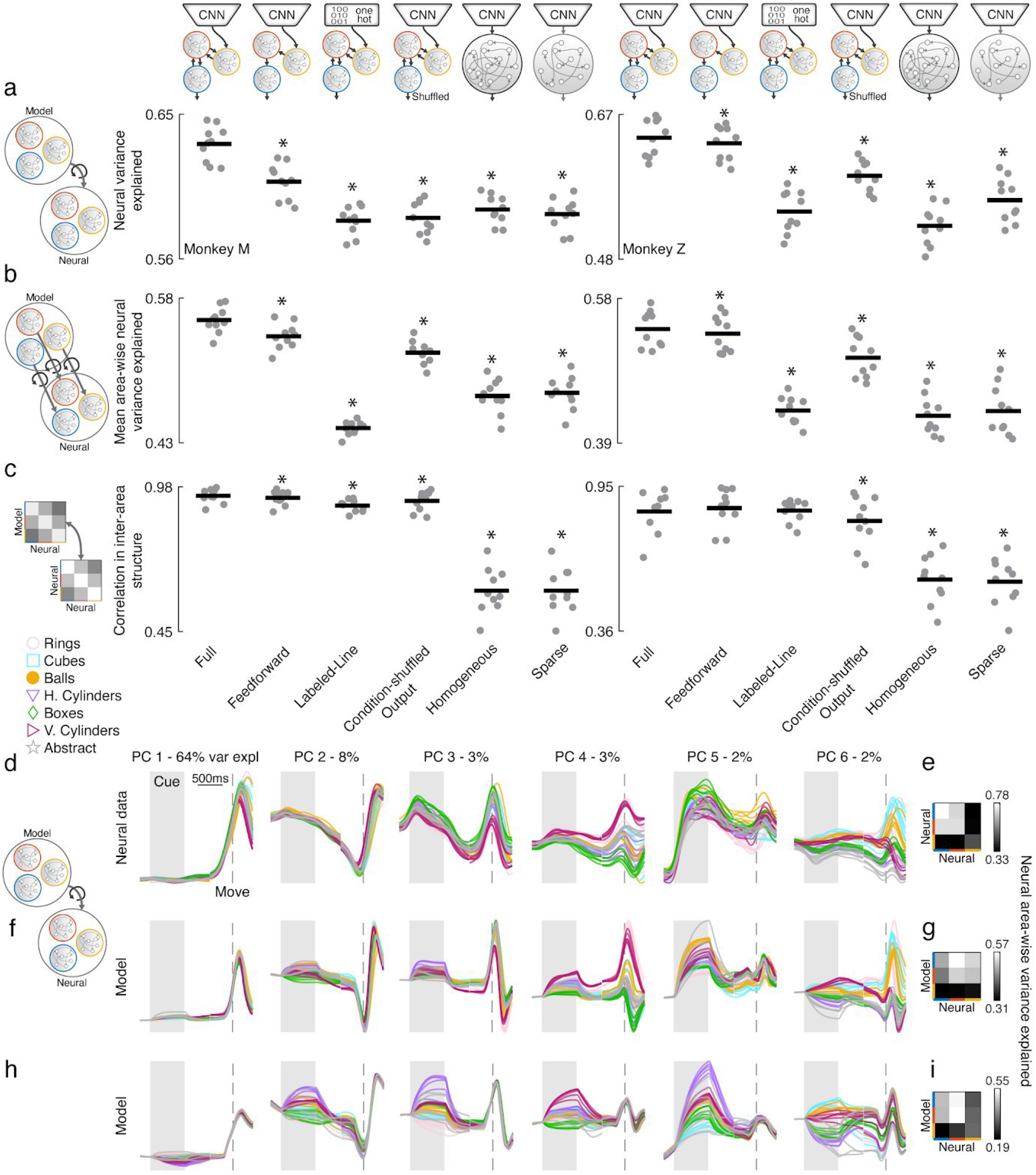
Modular RNN with visual feature input outperforms alternative models of neural data from the grasping circuit. (a) Average performance per recording session for the best set of regularization parameters for each architecture (averaged over 5 runs) for each of the three proposed metrics, procrustes across regions (a), average area-wise procrustes (b), and the match in inter-area structure between model and neural data (c). Horizontal bars represent the mean, and each dot represents a single session. Stars indicate significant difference as compared to Full model (paired t-test, p < 0.01). We tested 5 alternative models in addition to the Full model: 2) mRNN model with only feedforward connections between modules, 3) mRNN model receiving a labeled-line input (one-hot), where each condition is represented by a separate input dimension, 4) mRNN model with output conditions shuffled (objects reassigned), 5) homogeneous, fully-connected module, or 6) a single, sparsely-connected module with sparsity matching the Full model. (d,f,h) Procrustes analysis comparing the dynamics of two exemplar models to neural data across all brain regions (session M2). Fits of the simulated to neural data were projected onto the first 6 PCs defined on the neural data for visualization purposes only, and percentages show variance explained in biological data per PC. (e,g,i) Area-wise procrustes was performed between each brain region and itself, or between each module and brain region. (f) Exemplar model with the parameters (Homogeneous model, ReTanh activation function, L2 rate regularization - 1e-1, L2 weight regularization - 1e-5, inter-module sparsity - 0.1). (h) Exemplar model with the parameters (Condition-shuffled output model, ReTanh activation function, rate regularization - 1e-1, weight regularization - 1e-5, inter-module sparsity - 0.1). For d,f,h, the multiple traces for each type of object represent the different sizes within a turntable.

### Models generate grasping movements using simple dynamical rules

Earlier in Figure 2, we quantified how the representation of the task conditions within the grasping circuit shifted between visual cue onset and movement onset, posing the question of *how* and *why* this shift in representation occurs. Since our models are fully observable, we were able to probe our networks for explanations of the computations necessary for the task using fixed point analysis (Chaisangmongkon et al., 2017; Sussillo and Barak, 2013; Sussillo et al., 2015). In fixed point analysis, we perform an optimization to look for equilibrium points in the activity of the network, linearize the dynamics around these points, and interpret the properties of these linear dynamics (Methods). Whenever the input to a system changes, the fixed point structure changes. Therefore, we opted to perform this analysis jointly across all modules of the network during the 3 time epochs within which inputs were stable.

During the cue period, we found for each presented object a single fixed point that activity was attracted towards and was determined by the object presented (Fig. 6a - top), while similar activity was observed in the neural data (Fig. 6a - middle). These fixed points tended to be stable, with the majority of modes attracting activity to a point, organizing activity based on each specific object. Replacing the full model with the linearized system around each fixed point revealed that these fixed points were sufficient to capture the majority of the dynamics of the networks (Fig. 6a - bottom, linearized system explaining 97% of the variance in the full model). Interestingly, during the memory period we found a single fixed point (Fig. 6b - top) that maintained activity depending on the condition while also shifting the relative positions of conditions somewhat, similar to the neural data (Fig. 6b - middle), and was well captured by the linearized system (Fig. 6c - bottom, linearized system explaining 93% of the variance in the full model). Finally, during the movement we found a single fixed point with one or more modes of oscillation (Fig 6c - top), that rotated activity following these oscillatory modes and dependant on the starting position as determined by the memory period and the release of the hold signal. Neural activity showed a very similar pattern (Fig. 6c - middle) and once again the linearized system captured most of the dynamics (Fig. 6c - bottom, linearized system explaining 87% of the variance in the full model).

**Fig. 6.**
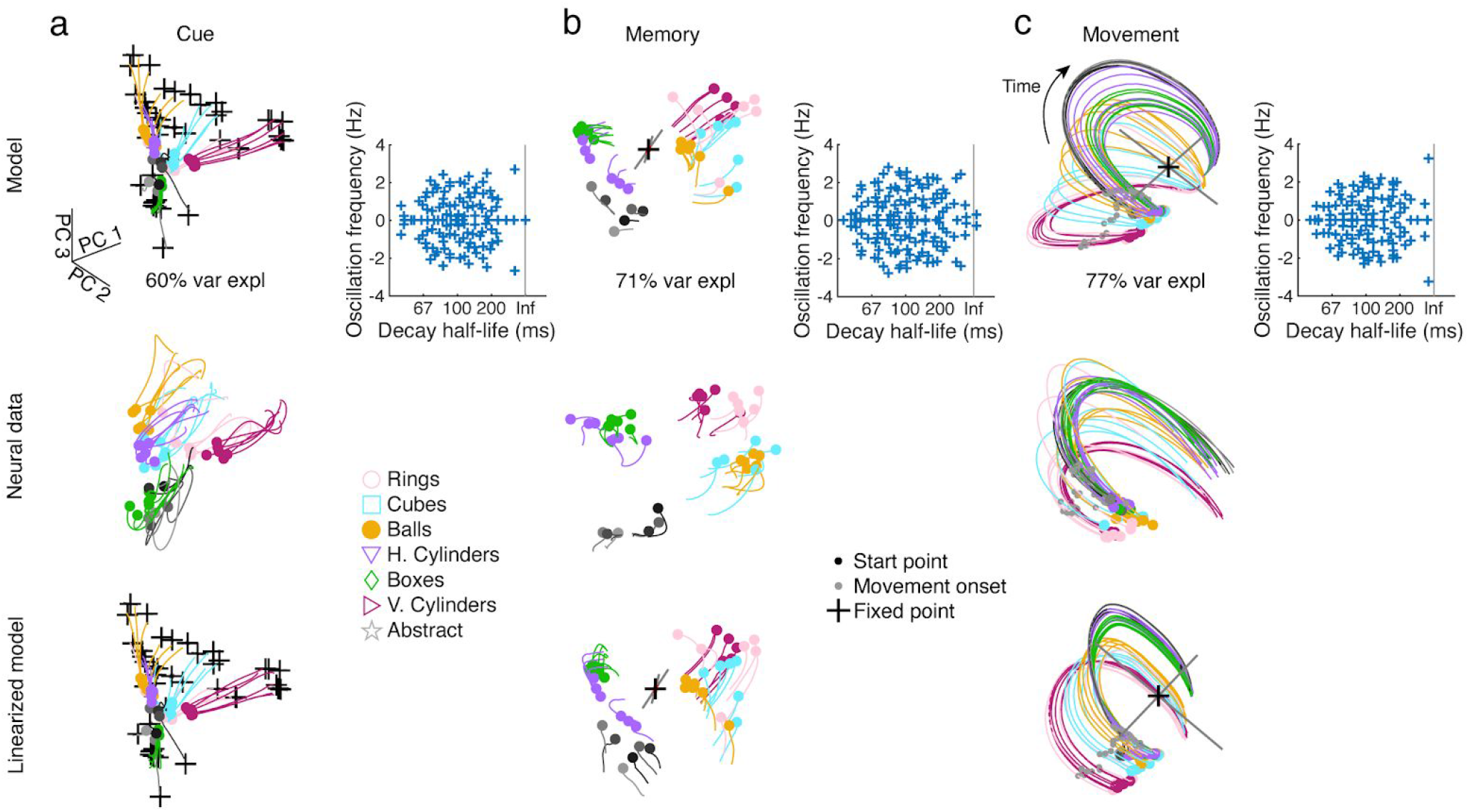
Fixed point analysis reveals simple computational strategies. (a - top) Fixed points of an exemplar Full mRNN model (same parameters as Fig. 3) during the portion of the cue period where inputs were stable (cue onset + 50 ms to cue offset) plotted in the first three PCs alongside trial-averaged activity, showing many fixed points, each corresponding to a different condition. Inset shows the complex eigenvalue spectrum of the linearized system around a representative fixed point. (a - middle) Neural data from an example session (M6) over the same time period, projected into the activity of the model using procrustes. (a - bottom) Replacing the full non-linear model with the linearized system around each fixed point yields similar trajectories. (b) Same as a for the memory period (cue offset + 50 ms to cue offset + 500 ms), showing a single fixed point in the model. The eigenvectors of the two largest eigenvalues are plotted (gray) for each fixed point, scaled by the magnitude of the eigenvalue. (c) Same as a for the movement period (150 ms before movement onset to 400 ms after movement onset), showing a single fixed point with an oscillatory mode.

We repeated this analysis for every network architecture, but found only minor differences between fixed point topologies employed by the different models, suggesting that this solution (1) is a parsimonious solution regardless of network architecture, and that (2) this solution is primarily a function of the task goal being optimized and not the architecture. Together, this analysis reveals that networks used simple computational strategies to reorganize and subsequently unfold activity during movement to generate the required muscle kinematics.

### Targeted lesioning of modules in rate regularized mRNNs reproduces behavioral deficits observed in the biological circuit

When M1 is lesioned, macaque monkeys lose the ability to shape the digits of the hand (Murata et al., 2008; Passingham et al., 1983), movements become smaller in amplitude, and the precise timing of muscle control is severely affected (Hoffman and Strick, 1995). On the other hand, reversible inactivations of either F5 or AIP cause monkeys to generate inappropriate hand shapes for the object they are grasping, mostly maintaining their ability to grasp after making contact with the object (Fogassi et al., 2001; Gallese et al., 1994).

Given the success of the mRNN models in explaining neural dynamics and inter-area differences in the grasping circuit, we were curious what behavioral deficits would be predicted from targeted lesioning of each module by silencing random subsets of neurons. We performed silencing with 240 variations, with each experiment additionally repeated 100 times with a different subset of neurons (Fig. 7a). When considering normalized kinematic error (Fig. 7b), it was clear that 1) networks with large amounts of rate regularization performed best, and 2) network performance clustered by module silenced, with best to worst ordered from input to output, respectively. To further examine the behavior for networks with the highest levels of rate regularization, we examined 4 behavioral metrics.

**Fig. 7.**
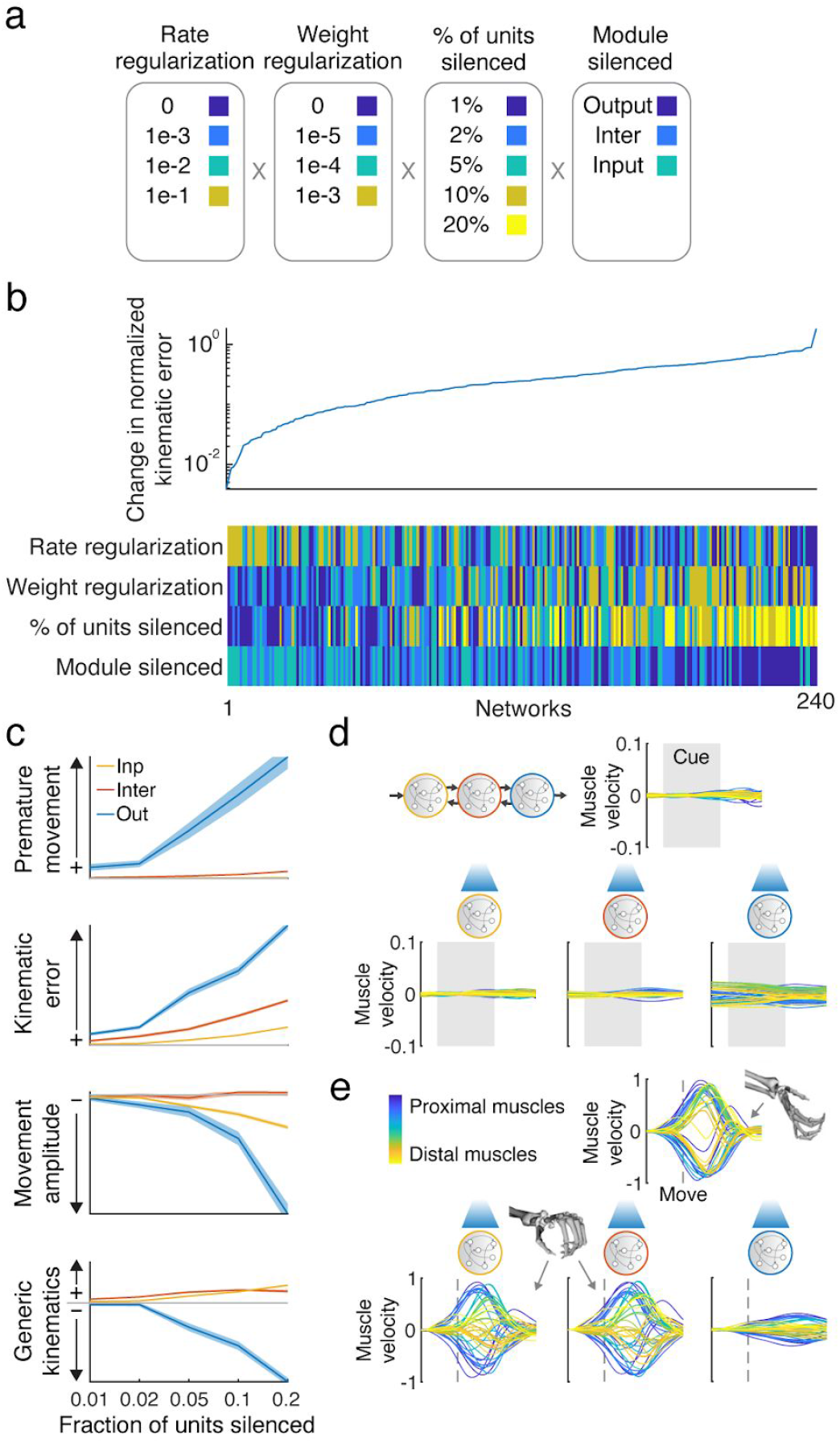
Targeted lesioning of rate regularized modular networks produces unique behavioral deficits. (a) Activity within 240 networks were artificially silenced, varying the rate and weight regularizations, percent of units silenced (repeated 100 times with random units), and the module being silenced. (b) Average change in normalized kinematic error after silencing as compared to normal operation. (c) Changes in network behavior as a function of module and number of units silenced for an exemplar network with high robustness to silencing (i.e. high rate regularization). Premature movement refers to the change in variance of output behavior before movement was initiated. Kinematic error refers to the change in normalized kinematic error during the movement period. Movement amplitude refers to the change in absolute movement activity. Generic kinematics refers to the change in normalized error in output behavior as compared to the mean kinematic behavior across all conditions. Shaded error bars represent SEM over 100 lesion repetitions. (d) Example network output before movement where 20% of the units in each module were silenced (example condition: mid-sized cylinder). (e) Example network output during the movement without silencing (top; mid-sized cylinder) and when 20% of the units in each module were silenced (bottom), showing that hand shape was not matched to the object (mid-sized cylinder) when the input or intermediate module were silenced (similar to the grip for the small cube or ball) while being degraded by silencing of the output module.

Silencing the output module increased the amount of premature movement activity, while this was not the case for the other modules (Fig. 7c-d). The effect of silencing on kinematic error was graded by module, having the largest effect on the output module and least on the input module. Conversely, movement amplitude was attenuated when the output module was silenced, but less so when the other modules were silenced. Finally, while increasing the number of units silenced in the output module led to degraded behavior that lacked the spatiotemporal dynamics necessary for the task. In contrast, performing the same lesioning to the intermediate and input modules did not degrade behavior, but rather produced hand shapes that were not appropriate for the target object (Fig. 7c,e). In general, behavior became more generic, looking more like the mean kinematics across conditions, in this case most similar to the hand shape required for a small cube or ball. Similar results were obtained using rate regularized feedforward mRNNs. However, unregularized mRNN models did not show these unique behavioral deficits.

Interestingly, silencing all inputs to the output module from the intermediate module immediately after releasing the hold signal produced only minor deficits in the firing rate regularized mRNN (~10% normalized error), while catastrophically eliminating behavior in the non-regularized networks. This result suggests that the output module (M1) may be able to produce the required kinematics semi-autonomously during movement when firing rates are encouraged to be small, a hypothesis that has not been tested in the biological circuit.

Overall, these results show that targeted lesioning of modules with rate regularized mRNNs resembled behavioral deficits observed when lesioning the biological circuit, suggesting that our modular RNN may be able to explain specific motor deficits observed in a number of previous studies and that firing rate minimization may be an organizational principle across cortical regions.

## Discussion

Recurrent neural networks are powerful tools for generating complex temporal dynamics. In this work, we demonstrated that modular recurrent neural networks (mRNNs) trained to complete a complex behavioral task can strongly resemble the processing pipeline for grasping, the inter-area differences observed in the brain, as well as recapitulate some of the behavioral deficits caused by lesioning.

These mRNNs took in pixel data through a CNN and transformed them into the temporal muscle kinematics necessary to grasp various objects. Importantly, no neural data were involved in the model training procedure. Visual features of objects, as extracted by a convolutional neural network (CNN) trained to identify objects, provided inputs necessary to complete the task, and fit neural data better than alternative models (Figures 2 and 5), including networks with a simple labeled-line code or without modularity. Our results connect the many works on neural networks for object identity in the ventral stream (Bashivan et al., 2019; Cadieu et al., 2014; Yamins and DiCarlo, 2016) to grasp movement generation by showing that the features extracted by such networks are useful for generating grasping movements to learned objects, and that modular network models are essential for understanding the dorsal and ventral streams. Additionally, we found that modular networks with an intermediate level of inter-area sparsity (10%) explained neural data best, paralleling the biological circuit (Markov et al., 2014).

The visual inputs to our model were supplied by a CNN that has been generally compared to the ventral stream. This stream has primarily been implicated in object identity processing, while the dorsal stream is largely implicated in spatial localization (Maunsell and Newsome, 1987). Why was this network able to perform so well, despite the fact that AIP lies along the dorsal stream? We propose three reasons. Firstly, AIP is involved in extracting shape information for grasping (Theys et al., 2015), a process which likely requires the extraction of similar features to those useful for determining object identity. Secondly, AIP is strongly connected to ventral areas in the inferotemporal cortex, including TEa/m (Borra et al., 2008; Webster et al., 1994), and areas in the temporal cortex essential for 3D shape perception (Verhoef et al., 2012). These areas are thought to interact during 3D object viewing (Janssen et al., 2018) and are possible routes by which AIP could receive object identity information from the ventral stream. Lastly, in this task objects are only in one spatial location, essentially eliminating the need for a ‘where’ code that differs between objects, a dominant feature of the dorsal stream.

We found that architectures containing feedback between modules outperformed architectures with only feedforward connections at explaining neural data. However, these differences were relatively small, despite the fact that strong feedback connections exist in the anatomical circuit. The likely reason for this is that the task modeled in the current study is very feedforward in nature. Once the monkeys were trained on all objects, the object presented to the monkey on any given trial uniquely determined the grasp plan required to lift the object. These feedback connections would likely come into play in different tasks which have rules that determine how an object should be grasped in a given context. The object identity information that is relayed to AIP from TEa/m is also communicated to ventrolateral prefrontal (VLPF) cortex areas 46v (Gerbella et al., 2013) and 12r (Borra et al., 2011), which relay back to F5 and AIP (Fagg and Arbib, 1998; Gerbella et al., 2011; Grafton, 2010). These provide an anatomical substrate for context-dependent motor planning in the AIP-F5 circuit, something not explored in the current study. Future experiments should investigate objects in various locations, with rules and context, and try to close the loop by looking at haptic feedback from S2 to AIP (Borra et al., 2008) and F5 (Gerbella et al., 2011).

Fixed point analysis revealed that models were governed by simple dynamical rules. The single fixed points during the cue separated and reorganized conditions. During the memory period, a single fixed point dynamically maintained the task conditions in the absence of task-related input, similar to models of working memory in LIP/PFC (Chaisangmongkon et al., 2017). The organization of task condition was appropriate to set the initial conditions required to generate oscillatory patterns for grasping movements, paralleling results in the arm area of motor cortex (Churchland et al., 2010, 2012; Michaels et al., 2016; Russo et al., 2018; Sussillo et al., 2015), and suggesting that the initiation of grasping movements can be understood very similarly to reaching movements under the dynamical systems perspective (Shenoy et al., 2013). The amount of reorganization between cue presentation and movement initiation was more pronounced in the model than in the neural data, likely due to the fact that turntables were only changed periodically in the experiment, so neural activity at the beginning of trials was already organized by object type to some extent.

The fixed point topology of networks with differing architectures tended to be similar, even in non-modular networks. Previous work has shown that this is often the case in RNNs of differing architectures trained to solve the same task (Maheswaranathan et al., 2019). However, this does not preclude different predictions of these networks. For example, we found that modular networks regularized by firing rate predicted the behavioral deficits resulting from lesions to these areas, while non-regularized networks, or networks without modularity, could not. These results suggest that selection of the task goal is vastly more important for the resulting solutions than the particular architecture. Future work should put heavy emphasis on testing the differential effects of task goal, network architecture, and optimization procedure (Richards et al., 2019).

The density of long-distance connectivity between areas is predicted to decrease with increasing brain size (Ringo, 1991), likely following organizational principles of cortical geometry and an exponential distance rule (Ercsey-Ravasz et al., 2013; Horvát et al., 2016). However, it is not clear what computational benefit is bestowed by increased sparsity in the cortical graph. Communication efficiency appears to be largely conserved between the monkey and the mouse, even though the cortical graph of the mouse is much more dense (Gămănuţ et al., 2018), while some work suggests that modular neural architectures support higher complexity in neural dynamics (Pinto et al., 2019; Sporns et al., 2000). Overall, the current work is consistent with the notion that connectivity between cortical areas in the macaque is intermediately sparse (Markov et al., 2014), while the computational benefits and potential pitfalls (Bullmore and Sporns, 2012) of such connectivity requires further study.

Our lesion results provide intriguing evidence of how different behavioral deficits may emerge from lesions to different cortical regions responsible for grasping behavior. One aspect we do not address is potential differences that may emerge in reaching vs. grasping depending on the cortical region lesioned, since the reaching behavior in our study was almost identical across conditions. While dexterous movements suffer greatly from lesioning of descending motor pathways, gross movements such as reaching may be less affected (Lawrence and Kuypers, 1968a, 1968b; Lemon, 2008), potentially because of a larger subcortical contribution (Lemke et al., 2019). Future studies could address these differences by examining tasks where reach and grasp behavior are varied in tandem (Lehmann and Scherberger, 2013).

Our lesion results suggest that motor cortex could potentially act as an autonomous dynamical system during early movement, a question not resolved by previous work in reaching (Churchland et al., 2012). However, it has recently been shown in mice that the motor commands necessary for dextrous control cannot proceed without continuous input from the thalamus (Sauerbrei et al., 2020). One possibility is that while motor cortex in primates may be able to act autonomously during early movement after receiving a go signal, activity may shift to being more input driven once sensory feedback related to the movement arrives in motor cortex, a critical question to be tested in future work. Furthermore, these conflicting results reinforce the need for future models that consider sensory feedback as well as subcortical dynamics. Overall, this work builds on many years of work on goal-driven modeling, dynamical systems, and deep neural networks in the visual and motor systems to present a unified view of grasping from pixels to muscles.

## Methods

### Animal training and experimental setup

Experimental design has previously been described in detail (Schaffelhofer and Scherberger, 2016; Schaffelhofer et al., 2015a). Briefly, two rhesus monkeys (Macaca mulatta) participated in this study (monkey Z: female, 7.0 kg; monkey M: male, 10.5 kg). Animal housing, care, and all experimental procedures were conducted in accordance with German and European laws governing animal welfare and were in agreement with the guidelines for the care and use of mammals in neuroscience and behavioral research (National Research Council et al., 2003).

We developed an experimental setup that allowed us to present a large number of graspable objects to the monkeys while monitoring their behavior, neural activity, and hand kinematics. During each recording session, monkeys grasped a total of 36–42 objects of equal weight that were placed on 8 interchangeable turntables (Figure 1c), presented in random order for each recording session. Objects were of different shapes and sizes including rings, cubes, spheres, horizontal cylinders, vertical cylinders, and bars. A special turntable held objects of abstract forms, which differed visually, but required almost identical hand configurations for grasping. A mixed turntable held objects of different shapes of average size, but this turntable was excluded from the analyses in this study, since all objects were present on other turntables. Monkeys were also trained on a grasping box that cued one of two grasping types, power or precision grip, but for simplicity this data was also not included in the current study.

### Task paradigm

Monkeys were trained to grasp 42 objects in a delayed grasp, lift, and hold task (Figure 1c,d). While sitting in the dark the monkeys could initiate a trial (self-paced) by placing their grasping hand (left hand in monkey Z, right hand in monkey M) onto a rest sensor that enabled a fixation LED close to the object. Looking at (fixating) this spot for a variable time activated a spot light that illuminated the graspable object. After the light was turned off the monkeys had to withhold movement execution until the fixation LED blinked for 100 ms. After this, the monkeys released the rest sensor, reached for and grasped the object and briefly lifted it up (500 ms). The monkeys had to fixate the LED throughout the task (max. deviation: ~5 deg of visual angle). All correctly executed trials were rewarded with a liquid reward (juice) and monkeys could initiate the next trial after a short delay. Error trials were immediately aborted without reward and excluded from the analysis.

### Kinematic recording

Finger, hand, and arm kinematics of the acting hand were tracked with an instrumented glove for small primates. Eight magnetic sensor coils (model WAVE, Northern Digital) were placed onto the fingernails, the hand’s dorsum as well as the wrist to compute the centers of 18 individual joints in 3D space, including thumb, digits, wrist, elbow and shoulder. The method and its underlying computational model have been described previously (Schaffelhofer and Scherberger, 2012). Recorded joint trajectories were then used to drive a 3D-musculoskeletal model (Holzbaur et al., 2005; Schaffelhofer et al., 2015b), which was adjusted to the specific anatomy of each monkey. The model was implemented in OpenSim (Delp et al., 2007) and allowed extracting a total of 27 DOF in joint angle space, and 50 DOF in muscle tendon length space. All extracted joint angles and muscle lengths were sampled at 100 Hz and low-pass filtered (2^nd^-order Butterworth filter, 3 Hz low-pass).

### Electrophysiological recordings

Single and multiunit activity was recorded simultaneously using floating microelectrode arrays (FMA, Microprobe Inc., Gaithersburg, MD, USA). In each monkey we recorded 192 channels from 6 individual arrays implanted into the cortical areas AIP, F5, and M1 (Figure 1b). In each array, the lengths of the electrodes increased towards the sulcus and ranged from 1.5 (1^st^ row) to 7.1 mm (4^th^ row). In area F5, one array was placed in the posterior bank of the inferior arcuate sulcus approximately targeting F5a (longer electrodes) and approaching the F5 convexity (F5c; shorter electrodes). The second and more dorsally located array was positioned to target F5p. In AIP, the arrays were implanted into the end of the posterior intraparietal sulcus at the level of area PF and more dorsally at the level of area PFG. In M1, both arrays were placed into the hand area of M1 into the anterior bank of the central sulcus at the level of the spur of the arcuate sulcus (Rathelot and Strick, 2009). Surgical procedures have been described previously (Schaffelhofer et al., 2015a). Neural activity was recorded at full bandwidth with a sampling frequency of 24 kHz and a resolution of 16 bits (model: RZ2 BioAmp Processor; Tucker Davis Technologies, FL, USA). Neural data was synchronously stored to disk together with the behavioural and kinematic data. Raw recordings were filtered offline (bandpass cutoff: 0.3–7 kHz) before spikes were detected (threshold: 3.5x std) and extracted. Spike sorting was processed in two steps: First, we applied super-paramagnetic clustering (Quiroga et al., 2004) and then revised the results by visual inspection using Offline Sorter (Plexon, TX, USA) to detect and remove neuronal drift and artefacts. No other pre-selection was applied and single and multiunit activity were analyzed together.

### Visual and muscle feature analysis

In order to model the visual features of the objects being presented in the grasping task, we generated simulated images from the monkey’s perspective (Fig. S1). We preprocessed and fed these images into a convolutional neural network (CNN), VGG (Simonyan and Zisserman, 2014), that used spatial convolution over pixels to determine the identity of objects in an image. VGG was pre-trained on ImageNet (Deng et al., 2009), a massive set of labeled images. We did not train VGG on our images.

To test how well features within the layers of VGG could explain neural activity during early presentation of the objects (early cue period, 400 ms), we first transformed the responses of each layer of VGG to the presentation of all objects into its first 20 principal components, which explained 91-99% of the signal variance. Next, we regressed the features of each layer onto the single trial spike counts during the cue period of each unit separately (Matlab function *fitrlinear*), using 10-fold cross-validation. All regressions had a standard L2 (ridge) penalty of λ = 1/*n*, where n was the number of in-fold observations. We then took the median r-value over all units within a recording session, and reported the mean of those values across recording sessions in Figure 2b. This method is meant to mimic the BrainScore metric used in previous works of the visual system (Schrimpf et al., 2018).

In order to make comparisons between regions, we must control for differences in recording quality. Therefore, we calculated the internal consistency of each area (Schrimpf et al., 2018), which provides a measure of reliability across trials within a given condition. To calculate internal consistency, trials within each condition were split in half, forming two sets of trials. These sets were correlated with each other for each unit separately, and the resulting r-value was Spearman-Brown corrected to account for the halving of sampling size. We took the median of this distribution and repeated the above analysis 1000 times with different random partitions of trials and took the average over repetitions. Finally, the results of the regression in Figure 2b were normalized on a per-neuron basis by the internal consistency in Figure 2b-inset.

For the analysis of muscle kinematics in Figure 2d we performed the same regression analysis using the average muscle velocity of all 50 muscles during movement initiation (200 ms before - 200 ms after movement onset) to predict neural spike count during the same time period.

### Modular recurrent neural network

In order to model the planning and execution of a grasping task, we implemented the dynamical system, 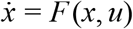, using a standard continuous RNN equation of the form

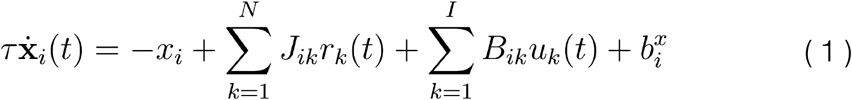

where the network has *N* units and *I* inputs, *x* are the activations and *r* the firing rates in the network, which were related to the activations by either the rectified hyperbolic tangent function (ReTanh), such that *r* = {0, *x* < 0; *tanh*(*x*), *x*≥0}, or a rectified linear function (ReLU), such that *r* = {0, *x* < 0; *x*, *x*≥0} . The units in the network interact using the synaptic weight matrix, *J*. The inputs are described by *u* and enter the system by input weights, *B* . Each unit has an offset bias, 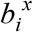. The time integration constant of the network is τ.

For all simulations *N* was fixed at 300, where each module contained 100 units (*N*_*m*_). The inputs were a condition-independent hold signal that was released 200 ms before movement onset and was sent to all modules, and a 20-dimensional signal representing the visual features of the current visual stimulus that was sent only to the input module. The elements of *B* were initialized to have zero mean (normally distributed values with 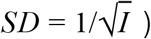. The elements of *J* were initialized to have zero mean (normally distributed values with 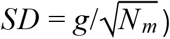 within each module, normally distributed with 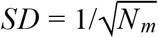 between each connected module, and zero for all other connections. The synaptic scaling factor, *g*, was set at 1.2 following previous work (Sussillo and Abbott, 2009). We used a fixed time constant of 100 ms for τ, with Euler integration every 10 ms. In addition, the sparsity of the connectivity between (but not within) modules was manipulated for the results in Figure 4.

The network was required to generate average muscle velocities in 50 dimensions until 400 ms after movement onset, where movement onset was determined by a threshold crossing in elbow position that approximately corresponded to the hand lifting off the handrest. The output of the network was defined as a linear readout of the output module

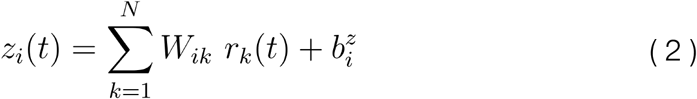

where *z* represents the 50-dimensional muscle velocity signal and is a linear combination of the internal firing rates using weight matrix *W*, which was initialized with near zero entries, and 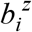, which is a bias term for each output dimension.

All non-zero values of the input weights, *B*, internal connectivity, *J*, output weights, *W*, and all biases, were trained using Hessian free optimization (Martens and Sutskever, 2011) (code: https://github.com/JonathanAMichaels/hfopt-matlab) also utilized in previous work (Michaels et al., 2016; Sussillo et al., 2015). The error function used to optimize the network considered the squared error between the output of the linear readout and the desired muscle velocity profiles, *v*

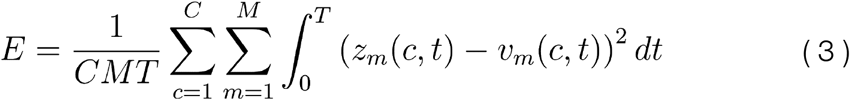

across all time, T, all muscles, M, and all conditions (trials), C, the networks were trained on. We report normalized error, which is the sum of the squared error from Eq. 3 over all times, dimensions, and trials, divided by the total variance of the target signal. In addition to the above error signal, we also implemented two commonly used regularizations. The penalties were a standard L2 cost on the firing rates, *R*_*FR*_, to keep units from saturating, and a standard L2 cost on the input and output weights, *R*_*IO*_. Therefore, the total error function minimized during training was

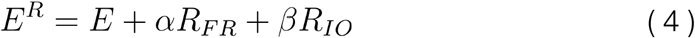

The hyper-parameter values, α and β, were varied systematically in the results of Figures 4–7 to test the effect of each regularization. The two regularizations were defined as

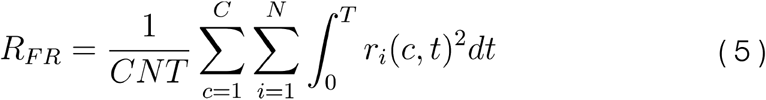

and

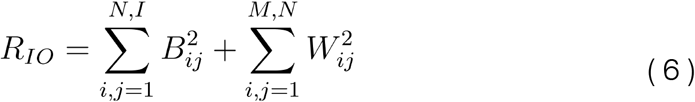

Similar to previous work, we opted not to model any feedback, since the goal of the study was to illustrate the main points parsimoniously and without relying on confronting the issue of what kind of feedback is most biologically plausible in such a network. All networks were trained until the change in objective from one iteration to the next fell below 1e-6.

### Assessing similarity of biological and neural data with procrustes

Imagine we have two shapes in front of us, for example a square and a triangle, consisting of the set of two dimensional points that make up each shape (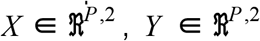, P - points). If we would like to see if it’s possible to overlap the triangle on the square *without distorting the overall shapes*, procrustes analysis provides a method for finding the optimal rotation that aligns them in arbitrary dimensionality. First, any differences in position and scale are removed by centering each object on a common coordinate system and by scaling all points by the Frobenius norm (yielding 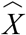 and 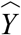. Then, the optimal rotation matrix, *R*, to align the triangle with the square can be found as follows: 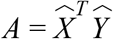, where the singular value decomposition *A* = *U* Σ*V*^*T*^ yields *R* = *UV*^*T*^ subject to *det*(*R*) = 1, making it a special orthogonal matrix. We can then quantify the success of the rotation by calculating 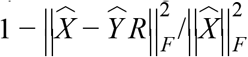, which would yield 1 for a perfect fit. This method is ideal for comparing simulated and biological data, where the square represents the neural activity of simulated data and the triangle biological data (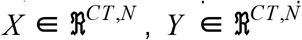, C - conditions, T - time points, N - neurons).

### Assessing inter-area structure with procrustes

To determine the estimated relationship between brain regions in the recorded data, we resampled trials (with replacement, equal to the number of recorded trials) for each recording session separately, condition-averaged the data, then performed pairwise procrustes between each pair of brain regions using resampled data. This procedure was repeated 100 times per recording session, and the average result was used to produce the 3 by 3 similarity matrices used to evaluate how well the modules matched the brain regions they were predicted to explain (e.g. Fig 3f,h).

### Fixed points

To extract simple rules behind the computations of our simulations, we searched for fixed points using standard nonlinear dynamical systems methods combined with linear stability analysis, as has been described in detail previously (Sussillo and Barak, 2013; Sussillo et al., 2015). We searched for a set of points in the high-dimensional state space, {*x*^1*^, *x*^2*^, …}, where the dynamics described in Eq. 1 are at an equilibrium, 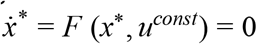, for a given constant input. For some volume around these points, Eq. 1 can be replaced by a linear dynamical system, 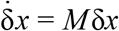, with δ*x* = *x* − *x*\* and 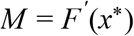, by definition. These points are considered fixed points if their speed is very slow relative to the speed of the network during normal operation (>1000 times slower for most of our results).

For each time epoch of interest, we repeated the optimization many times and randomly sampled the optimization starting point from activity the networks visited during normal operation, yielding a single fixed point for each condition during the cue period, a single fixed point during the memory period, and a single fixed point during the movement period. In some cases, tight clusters of fixed points (2-3) with similar properties were considered a single fixed point.

## Acknowledgements

We would like to thank N Bobb, L Burchardt, M Dörge, R Lbik, K Menz, and M Sartori for assistance, and B Dann, JA Gallego, M Perich, F Willett, and S Vyas for constructive discussions and helpful comments on an earlier version of this manuscript.

## Author Contributions

JAM, SS, and HS designed and carried out experiments; AAT, JAM, and SS analyzed data; JAM wrote the manuscript; All authors edited the manuscript.

## Supplemental Materials

**Fig. S1.**
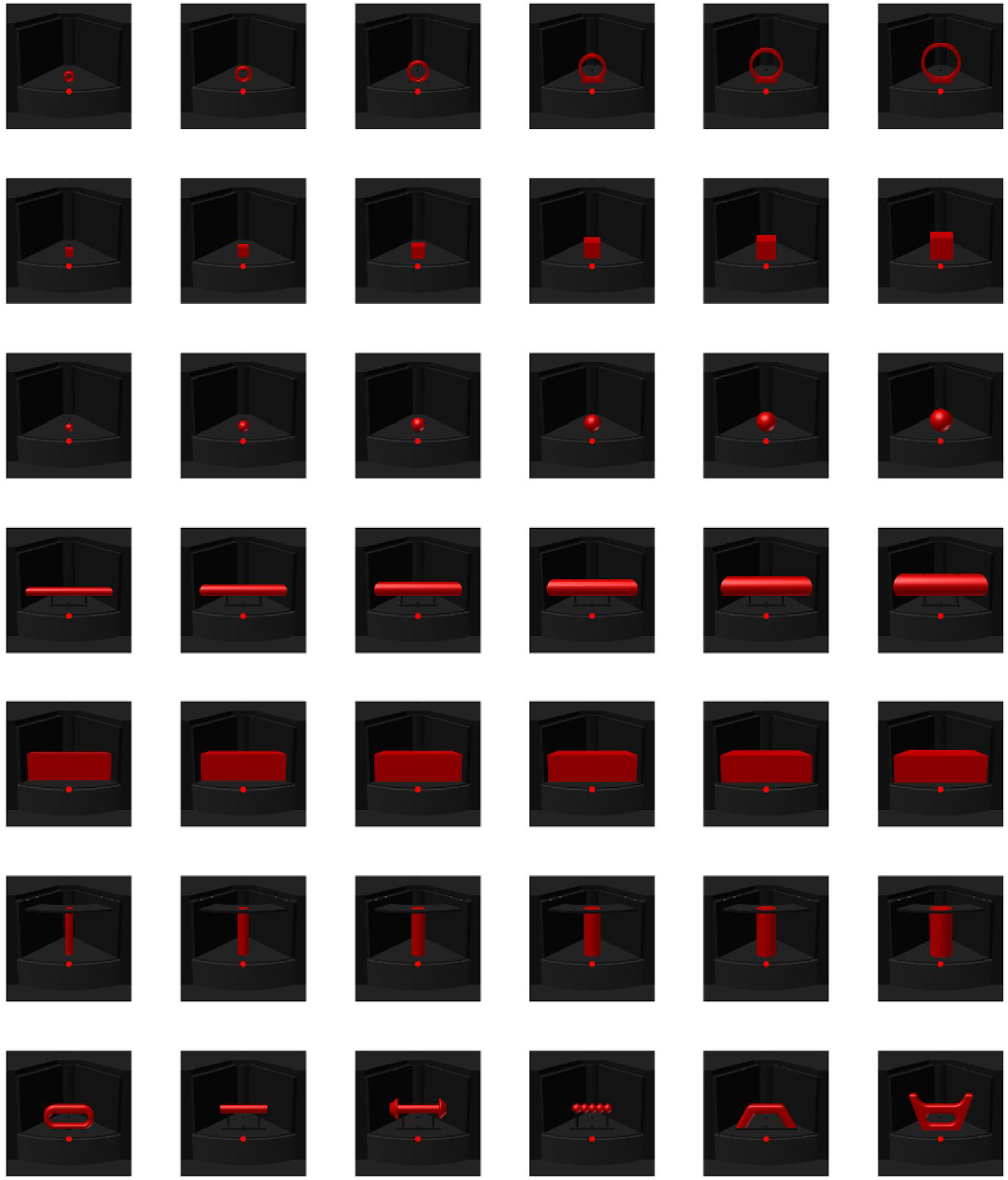
Simulated monkey view of objects used as input for VGG. Physical objects were CNC manufactured based on mesh models. Red fixation point was added in the approximate location that it was presented to the animals. All input images were RGB and 227×227 pixels in size.

**Fig. S2.**
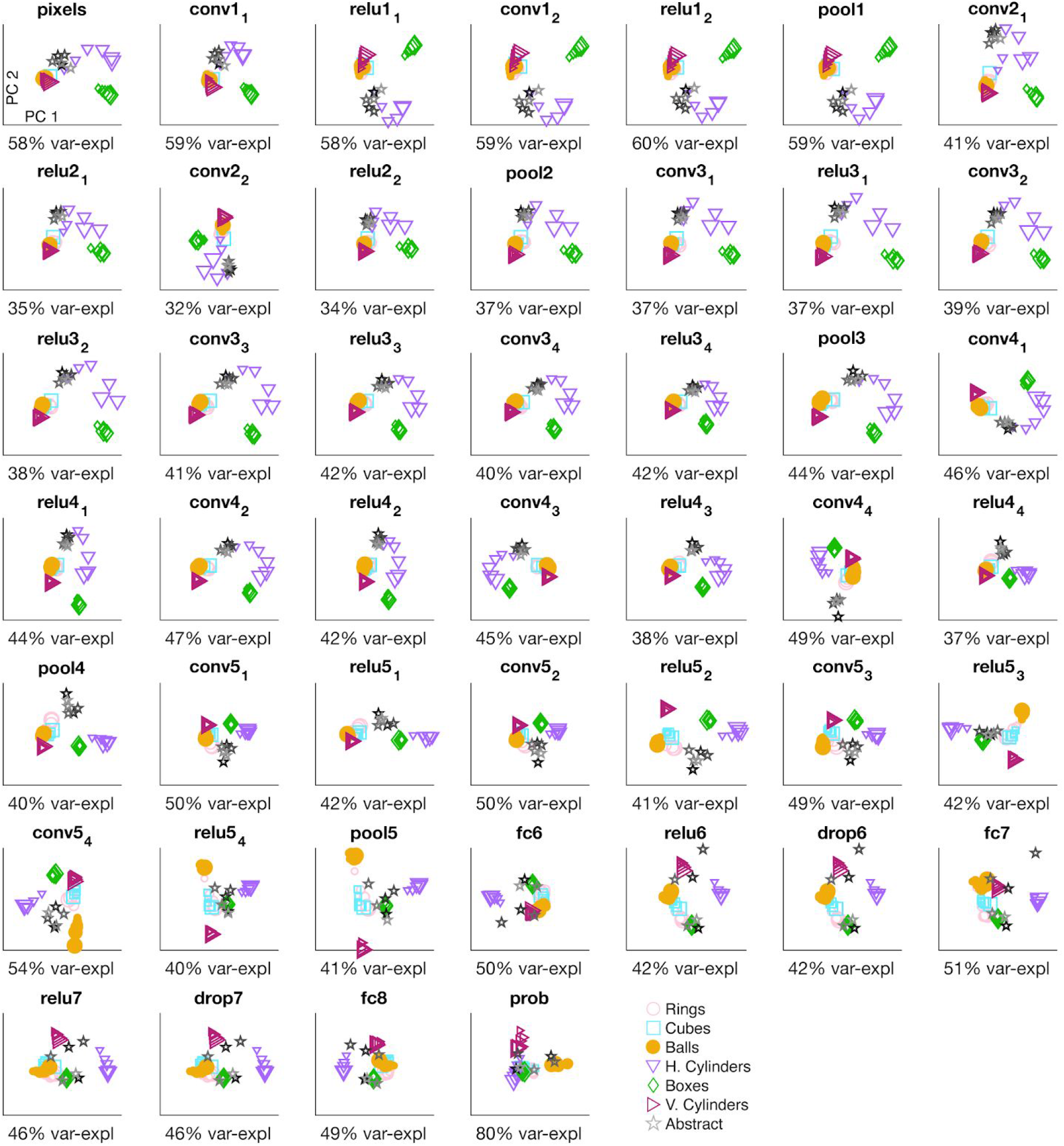
Feature representation in VGG. Representation of the features of all conditions in the first two principal components of each layer in VGG with corresponding variance explained.

**Fig. S3.**
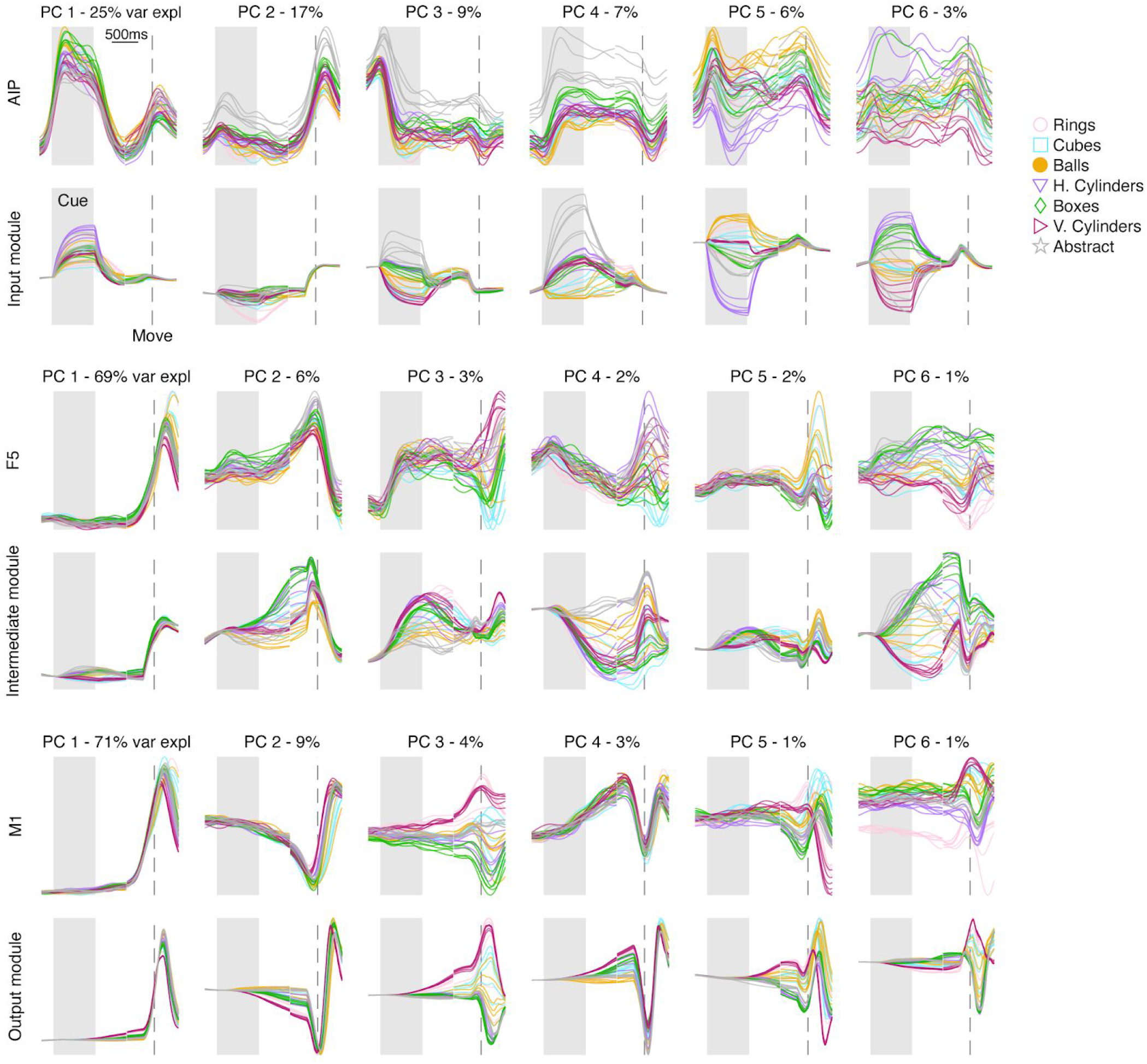
Area-wise ability of mRNN model to explain neural data in AIP, F5, and M1. Procrustes analysis comparing the dynamics of each module in an example model (Fig. 3) to the neural data of each brain region (session M2). Fits of the simulated to neural data were projected onto the first 6 PCs defined on the neural data for visualization purposes only, and percentages show variance explained in biological data per PC. The multiple traces for each type of object represent the different sizes within a turntable.

**Fig. S4.**
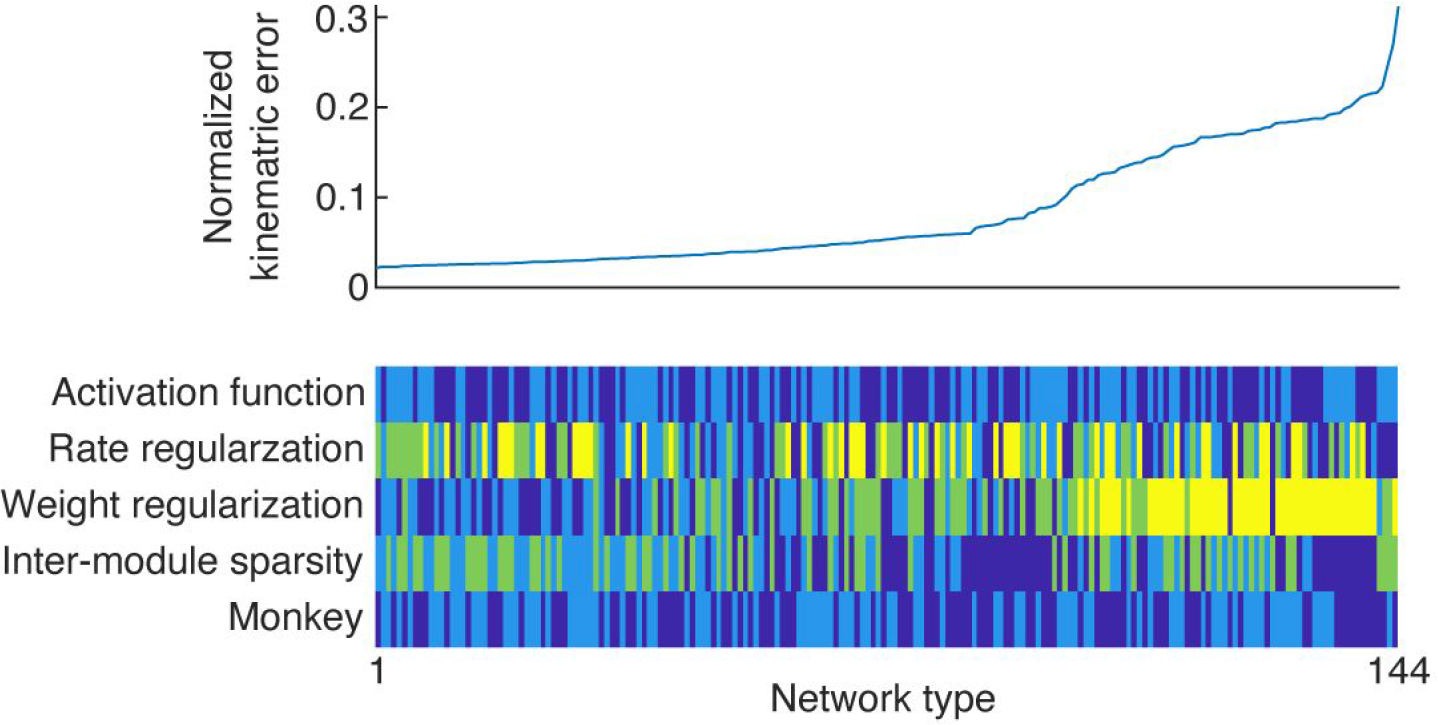
Normalized kinematic error across choice of network parameters. Average kinematic error for networks presented in Figure 4. For color code, see Fig. 4a.

